# Paralog-dependent Specialization of Paf1C subunit, Ctr9, for Sex Chromosome Gene Regulation and Male Germline Differentiation in Drosophila

**DOI:** 10.1101/2024.04.23.590698

**Authors:** Toshie Kai, Jinglan Zheng, Taichiro Iki

**Affiliations:** Laboratory of Germline Biology, Graduate School of Frontier Biosciences, Osaka University. Yamadaoka1-3, Suita, Osaka, Japan

**Author notes:** Correspondence: Taichiro Iki, Toshie Kai.

## Abstract

Testis-specific gene regulatory mechanisms govern the differentiation of germ cells into mature sperms. However, the molecular underpinnings are not fully elucidated. Here, we show the subunits forming Polymerase Associated Factor 1 Complex (Paf1C), a transcription regulator conserved across eukaryotes, have their individual paralogs predominantly expressed in *Drosophila* testes. One of these, namely, Ctr9 paralog enriched in testes (Ctr9t) was found to play a critical and nonredundant role in post-meiotic spermatid differentiation and male fertility in *D. melanogaster*. A proximity proteome analysis provides evidence that Ctr9t forms complexes with other paralog members. We show endogenous Ctr9t is expressed in germ cells at spermatocyte stages, accumulating in a distinct compartment within the nucleolus. There, Ctr9t co-localizes with Spermatocyte arrest (Sa), a testis-specific paralog of TATA-binding protein (TBP)-associated factor 8 (TAF8). We further demonstrate that *ctr9t* function is crucial for maintaining Sa in the nucleolus, but not *vice versa*. Transcriptome profiling reveals that Ctr9t acts as an activator for the set of male fertility genes on Y chromosome, but it also acts as a global repressor of X chromosome genes. Collectively, our results shed light on the nucleolus-associated, paralog-dependent regulation of gene expression from sex chromosomes, which ensures the terminal differentiation of male germ cells.

**Author summary:** A wide repertoire of genes takes a coordinated action during sperm formation. However, gene expression during spermatogenesis is controlled by heretofore unelucidated mechanisms. In this study, we show *Drosophila* expresses testis-specific paralogs of the individual subunits forming a conserved transcription regulator called Paf1C. We generated a loss of function allele of one of the paralog members which we named Ctr9t, and demonstrate that Ctr9t function is crucial for post-meiotic spermatid differentiation and male fertility. Therefore, the paralog has established a nonredundant role for spermatogenesis. Using newly generated antibodies, we show Ctr9t protein is expressed in germ cells at spermatocyte stages and, intriguingly, enriched in the nucleolus. There, Ctr9t interacts with Spermatocyte arrest (Sa), a key transcription factor essential for meiotic and post-meiotic differentiation processes. By further performing whole testis transcriptome analyses, we reveal that Ctr9t acts as a regulator of gene expression from sex chromosomes. Importantly, the activation of male fertility genes located on Y chromosome relies on Ctr9t function, which can explain the defects observed in *ctr9t* mutant germ cells. Altogether, we propose that Ctr9t acts as a key player in the spermatocyte nucleolus where it controls the expression of sex chromosome genes and ensures functional sperm formation.

## Introduction

Production of functional sperms is prerequisite for the life cycles of sexually reproducing organisms. Spermatogenesis follows a multistep differentiation program, involving changes in cell cycle dynamics, morphogenesis, organella organization, and chromatin remodelling, generating a highly polarized haploid gamete having condensed DNA and motile cilia [1–3]. A wide repertoire of genes takes a coordinated action during the complex processes, the expression of which is controlled by diverse transcriptional and posttranscriptional mechanisms [4–10]. Hence, spermatogenesis provides an attractive model for elucidating the principle of tissue/cell-specific gene regulation.

Gene expression programs in male germ cells require the functionally differentiated paralogs of general transcription regulators. In *Drosophila*, meiotic progression is preceded by transcriptional activation of an array of genes in pre-meiotic spermatocytes. This mass activation relies on two types of testis-specific regulatory apparatuses. One is an alternate of TFIID, formed by testis-specific TATA-binding protein-associated factors (tTAFs), Sa (dTAF8 paralog), Can (dTAF5 paralog), Rye (dTAF12 paralog), Mia (dTAF6 paralog), and Nht (dTAF4 paralog) [11–16]. The other module is meiosis arrest complex (tMAC), sharing a part of subunits with MMB/dREAM general repressor complex, but replacing some subunits with testis-specific homologs including Aly (Mip130 paralog), Tomb (Mip120 paralog), and Wuc (dLin52 paralog) [17–19]. tMAC and tTAF cooperatively act over a thousand of genes in spermatocytes, with the assist of Mediator complex and through the interdependency of subunit expression [20,21]. Generally, loss of either one of above paralogs leads to the characteristic meiotic arrest phenotype during spermatogenesis; the failure of entry to meiotic divisions or initiation of spermatid differentiation, demonstrating their functions are nonredundant and distinct from those of universally expressed counterparts. Beyond their paramount importance for meiosis, the paralog activities are supporting the post-meiotic terminal differentiation processes, given their responsibility for the expression of genes required at spermatid stages [13,19].

Transcription initiation is followed by the sequence of events including RNA polymerase II pausing, pause release, elongation, and termination, each of which is controlled by distinct sets of factors. One of crucial modules controlling post-initiation steps is Polymerase associated factor 1 complex (Paf1C), originally identified in yeast with the five subunits, Paf1, Ctr9, Cdc73, Rtf1, and Leo1 [22,23]. These subunits are widespread across eukaryotes with minor differences in their composition, their general functions as gene activators have been demonstrated in different biological contexts. In *Drosophila*, Paf1/Antimeros binding to Cdc73/Hyrax is recruited to transcriptionally active loci, being required for H3K4 trimethylation (H3K4me3) and transcriptional activation upon heat shock [24]. Rtf1 colocalizes with Paf1 in transcriptionally active loci, though it lacks stable binding to Paf1. The role of Rtf1 maintaining H3K4me3 levels has been shown in Notch signalling [25]. Gene activator function of Cdc73 has been demonstrated in Wnt signalling and hedgehog pathway [26,27]. The role of Ctr9 maintaining proper H3K4me3 levels has been demonstrated in nervous system and ovaries [28,29]. A recent study characterized Paf1 and Rtf1 as antagonist of PIWI-interacting (pi)RNA-mediated gene silencing in ovarian somatic cell culture [30], which is analogous to the fission yeast Paf1C that is opposing short interferring (si)RNA pathway [31]. Leo1 interacts with Myc and assists the binding to target gene promoters [32]. Paf1C subunit activities in testes remain uncertain.

Here, we identify and characterize the paralogs of Paf1C subunits that are preferentially expressed in *Drosophila* testes. We generated a loss of function allele of one of those, Ctr9 paralog, demonstrating that the mutant is defective in spermatid differentiation and functional sperm production. By further combining proximity proteome, immunohistochemistry, and transcriptome data, our study suggests that Ctr9t paralog localizes to the spermatocyte nucleolus, where it interacts with other factors to regulate gene expression from sex chromosomes for maintaining male fertility.

## Result

### Drosophila paralogs of Paf1C subunits predominantly expressed in testes

Paf1C is a complex conserved across eukaryotes, generally functioning as activator of transcription through the control of associating RNA Polymerase II and the modulation of chromatin states [22]. The core components, Paf1, Ctr9, Cdc73, Leo1, and the dissociable factor Rtf1, are widespread from yeast to humans (Figure 1AB). Transcripts encoding the individual subunits, *Paf1/Atms*, *Ctr9*, *Leo1/Atu*, and *Cdc73/Hyx*, are expressed in different tissues of *D. melanogaster*, including ovaries and testes (Figure 1C, see also FlyAtlas 2, https://motif.mvls.gla.ac.uk/FlyAtlas2/). Our BLAST search identified one paralog for each Paf1/Atms, Ctr9, Leo1/Atu, Cdc73/Hyx, respectively (Figure 1B). For Rtf1, three paralogs are found, of which tPlus3a and tPlus3B have been characterized previously as factors involved in spermatid differentiation (Figure S1A) [33]. Accumulating amino acid substitutions indicate the rapid evolution of paralogs compared to their individual counterparts (Figure 1B). These paralogs are widely conserved across *Drosophilidae*, however, for Paf1, Ctr9, and Leo1, we failed to identify their paralogs in *D. pseudoobscura* and the sympatric species, *D. persimilis* [34]. On the other hand, Cdc73 paralog can be found in the above two species, but not in *D. virilis*, *D. mojavensis*, and *D. grimshawi*.

**Figure 1.**
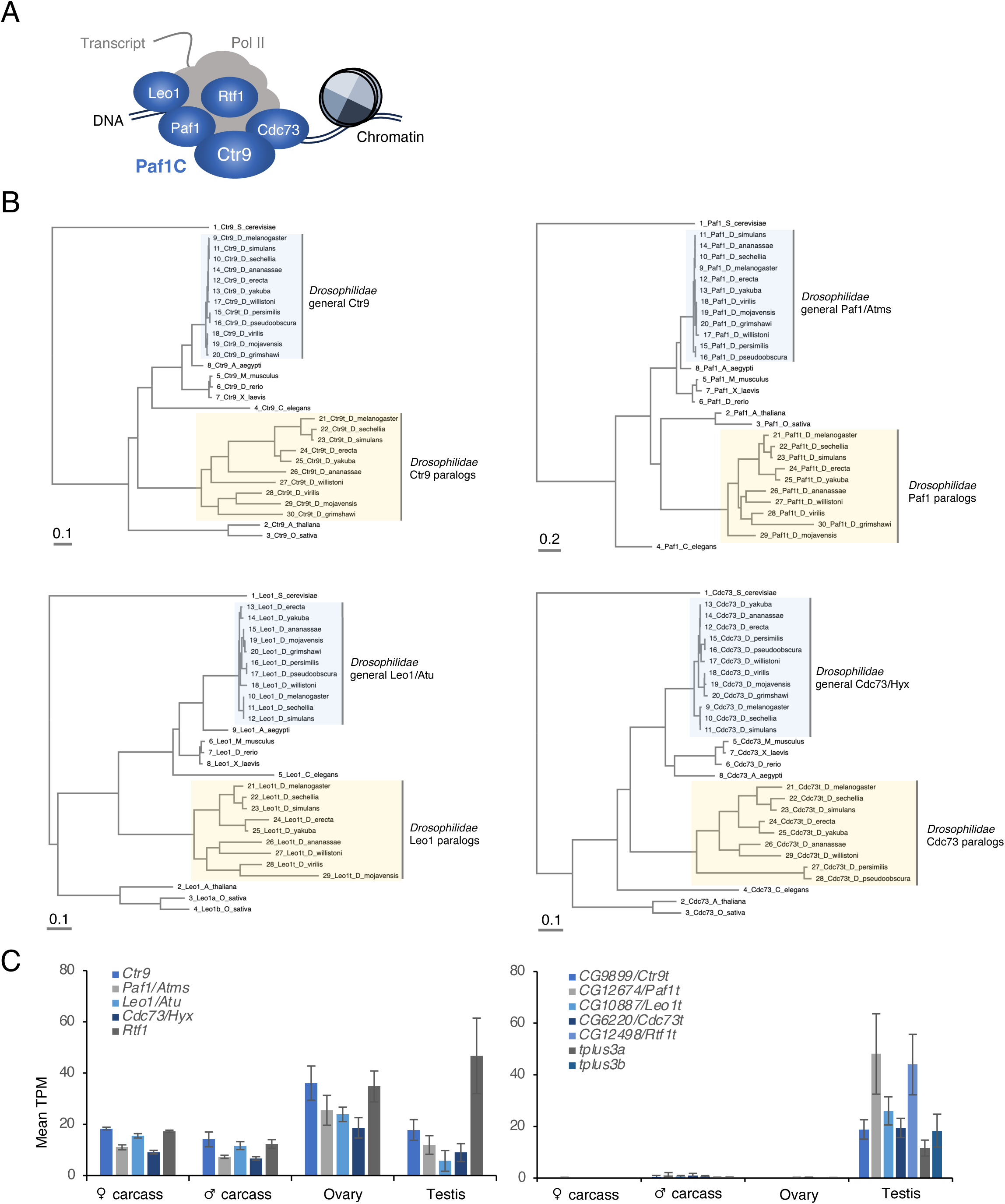
Paf1C subunit paralogs present in Drosophilidae. (A) Paf1 complex (Paf1C). Paf1C contains Ctr9, Paf1, Leo1, Cdc73 as core components. Rtf1 is known to be dissociable. Paf1C interacts with RNA pol II and control transciption-associated events. (B) Multiple alignment and phylogenetic analyses of individual Paf1C subunit proteins. Branch lengths measure the expected substitutions per site as indicated in the scale bar. (C) Expression patterns of genes encoding general Paf1C subunits (left) and their individual paralogs (right) in *D. melanogaster*. Source data: PRJEB22205.

The paralogs are predominantly expressed in the testes of *D. melanogaster*, which contrasts with their counterparts expressed universally (Figure 1C). Based on the characteristic expression patterns, the paralog genes are hereafter referred to as *ctr9 paralog enriched in testis (ctr9t), paf1 paralog enriched in testis (paf1t), leo1 paralog enriched in testis (leo1t), cdc73 paralog enriched in testis (cdc73t), and rtf1 paralog enriched in testis (Rtf1t)*, respectively.

### *ctr9t* is crucial for spermatid differentiation and male fertility

Given the preferential expression in testes (Figure 1), the paralogs of Paf1C subunits might play a role in spermatogenesis. To examine this possibility, a null allele of *ctr9t/CG9899* (*ctr9t^KO^)* was generated in *D. melanogaster* using CRISPR-Cas9-based genome editing [35]. Ctr9 homologs share the characteristic tandem array of tetratricopeptide repeat (TPR) motifs involved in Paf1C formation [22]. Introduced premature stop codon was expected to truncate the polypeptide upstream of most TPR motifs maintained by Ctr9t (Figure 2A S2A). In contrast to the null mutants of general *ctr9* that die before early larval stages [28,29], the homozygous *ctr9t* mutants (*ctr9t^KO/KO^*) were viable and grew normally to adulthood. Immunoblotting with newly generated anti-Ctr9t antibodies showed a distinct ∼100kDa band corresponding to the endogenous Ctr9t protein present in the lysates from *ctr9t^KO^/CyO* heterozygous control, while the band is absent in *ctr9t^KO/KO^* mutant testes (Figure 2B, S2B).

**Figure 2.**
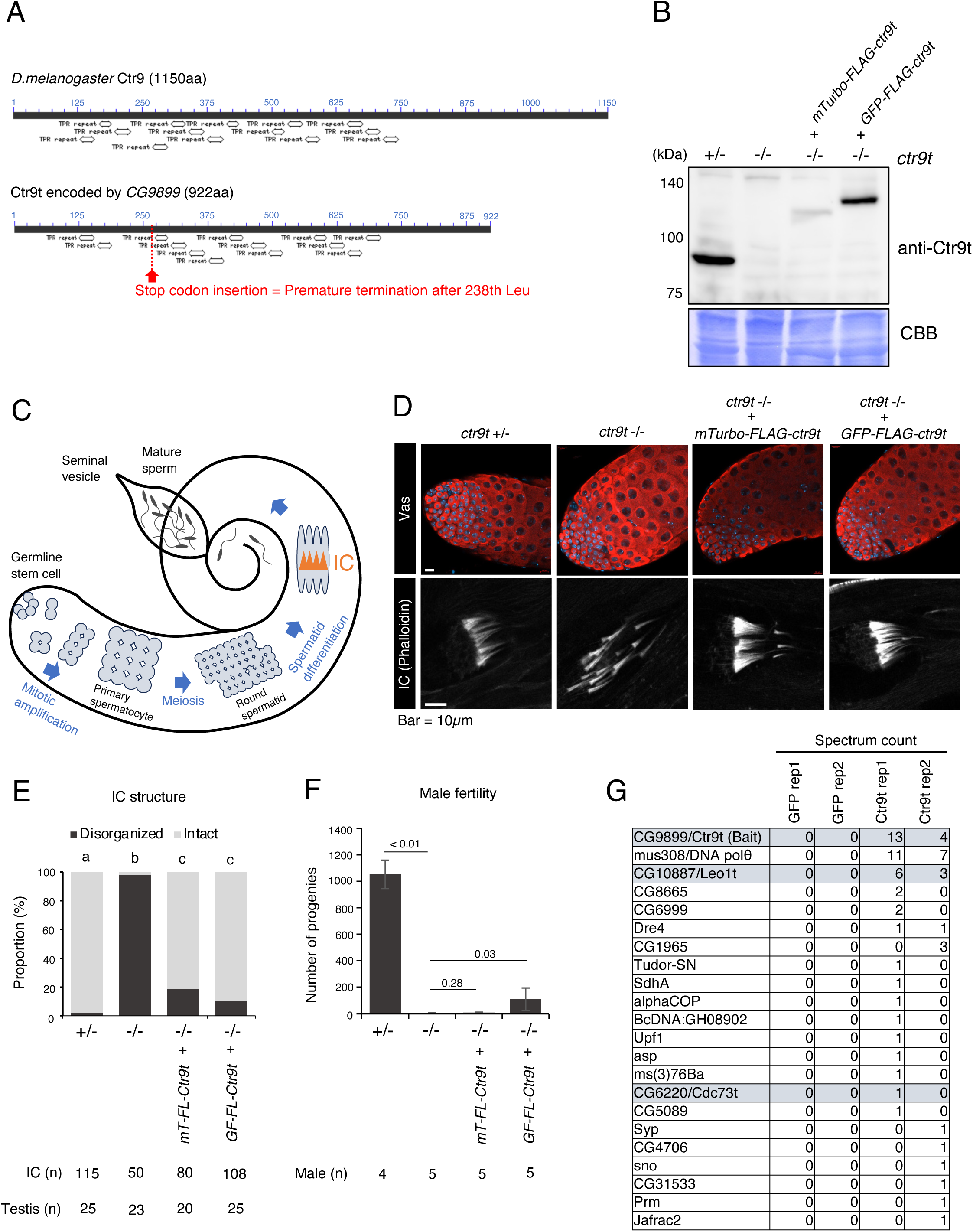
Genetic dissection of *ctr9t* function. (A) Polypeptide domain/motif analyses for Ctr9 and the paralogue Ctr9t. TPR; tetratricopeptide repeat. Images are taken from NCBI conserved domain platform. (B) Immunoblotting of testicular lysates with anti-Ctr9t antibody. +/-; *ctr9t*^KO/CyO^, -/-; *ctr9t*^KO/KO^. Transgenes are expressed under the control of *bam* promoter. Coomassie brilliant blue (CBB) staining serves as a protein loading control. (C) Spermatogenesis in Drosophila testis. Note that somatic cells associated with germ cells are not described. IC; individualization complex. (D) Top; germ cells (identified with Vas in red) in the apical end of testes. Bottom; IC (stained by phalloidin) in elongating spermatids. (E) IC organization. Proportion of disorganized ICs in elongating spermatids is shown. ICs containing more than five visibly dissociated actin corns are defined as “disorganized.” Different characters (a/b/c) indicate the significant difference (p < 0.01, Tukey test). (F) Male fertility. The number of progenies from single male are shown with p values (two-tailed unpaired t-test) were indicated. (G) Proximity proteome of Ctr9t. Identified paralogs of Paf1c subunits are highlighted in grey.

An adult testis maintains germ cells at distinct differentiation stages in a spatially ordered manner, from germline stem cells at the apical end to maturing sperms in the basal end (Figure 2C). *ctr9t* mutant testes showed no discernible defect in spermatogonia and early spermatocytes (Figure 2D upper panel). However, at spermatid stages, the characteristic F-actin investment called individualization complex (IC) became perturbed (Figure 2DE). In line with this, *ctr9t* mutant males were nearly sterile (Figure 2F). Similar phenotypes were observed also in *ctr9t^KO^/Df* transheterozygous mutants (Figure S2CD).

IC organization was largely recovered by Gal4/UASp-based expression of two different *ctr9t* transgenes (UASp-*mini(m)Turbo-FLAG-ctr9t* and UASp-*GFP-FLAG-ctr9t*) by germline-specific driver, *bam-Gal4*, indicating a critical role of *ctr9t* in germ cells (Figure 2DE) [36]. Consistently, male fertility was mildly recovered, but only with *GFP-FLAG-Ctr9t* (Figure 2F). Failure of recovery with *mTurbo-FLAG-Ctr9t* could be due to the unstable nature of the fusion proteins (Figure 2B). In addition, *bam*-Gal4/UASp system does not perfectly recapitulate the endogenous expression pattern (see relevant data in the following section). We also do not exclude a possibility that Ctr9t might function in somatic cells for other than IC formation. Overall, these results indiate that Ctr9t is functionally differentiated from general Ctr9 and plays a non-redundant role in spermatid differentiation and sperm production.

Like general Ctr9 forming Paf1C, Ctr9t proteins may be able to form Paf1C-like complexes with other paralog members in testicular germ cells. To examine this possibility, we analyzed the proximity proteome of Ctr9t using BioID/TurboID [37]. The mass spectrometry did not provide sufficient peptide signals, probably due to the weak expression of mTurbo-FLAG-Ctr9t (Figure 2B). Nonetheless, Leo1t and Cdc73t were found in the proximity of Ctr9t (Figure 2G). This result implies that Ctr9t can form a complex with other testis-specific paralog members in germ cells.

### Ctr9t localizes to a compartment of spermatocyte nucleolus

To understand the molecular feature of Ctr9t, we next examined the expression and subcellular localization of Ctr9t in testes. Immunostaining using anti-Ctr9t antibodies showed that Ctr9t was enriched in a distinct structure in germ cells at spermatocyte stages (Figure 3A). On the other hand, Ctr9t was undetectable in spermatogonia, which was confirmed by the absence of Ctr9t signal in *bag-of-marbles (bam)* mutant testes where germ cell differentiation was arrested early at mitotic amplification steps (Figure S3A). This structure enriched with Ctr9t was overlapped with a part of histone (H2Av-RFP), indicating that Ctr9t structure is formed inside the nucleus and associated with chromatin (Figure 3B). Ctr9t can partially overlap with DNA (DAPI). Confirming the immunostaining signals, a similar structure was observed by detecting the fluorescence from GFP-FLAG-Ctr9t (Figure S3B). Of note, the ectopic expression led to abnormal protein dispersion in the cytoplasm of spermatogonia and early spermatocyte cells.

**Figure 3.**
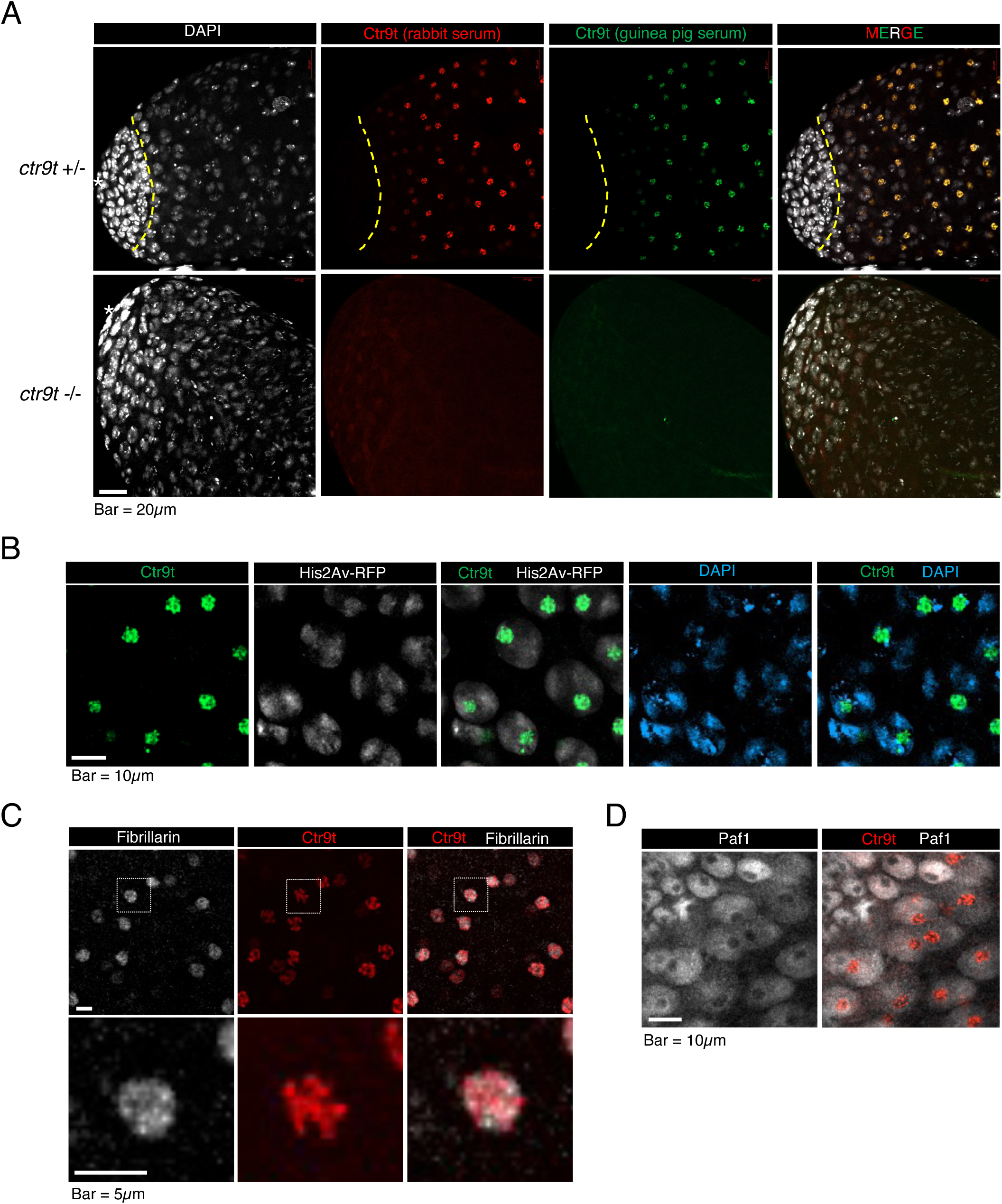
Ctr9t expression and localization in testes. (A) Testes immunostained for endogenous Ctr9t. Anti-Ctr9t antibodies raised in rabbit (red) and guinea pig (green) showed identical signals in spermatocyte nuclei. Dotted yellow line roughly delineates the boundary between spermatogonia and spermatocytes. (B) Ctr9t (green), histone (H2Av-RFP, white), and DNA (DAPI, blue) signals in spermatocyte nuclei. (C) Spermatocytes stained for Ctr9t (red) and a nucleolus marker, Fibrillarin (white). (D) Spermatogonia and spermatocytes immunostained for Paf1, a component of general Paf1C (grey), and Ctr9t (red).

Where within the nucleus does Ctr9t accumulate? One of the prominent subnuclear structure is the nucleolus where various biological processes including ribosome biogenesis take place [38]. Fibrillarin is a processing enzyme for ribosomal RNA, and thus serves as a useful nucleolus marker. We found Ctr9t and Fibrillarin occupied the same space in the nucleus, though the two signals did not match perfectly (Figure 3C). This result suggests that Ctr9t is localized to the nucleolus of spermatocytes and concentrated in a specific compartment. Paf1 and Cdc73, the subunits of general Paf1C, accumulated in the nucleoplasm, and did not co-localize with Ctr9t (Figure 3D, S3C). In addition, H3K4me3, a histone mark associated with general Paf1C was barely detectable in the nucleolar region (Figure S3D). These data support the notion that Ctr9t functions independently of general Paf1C.

### Ctr9t co-localizes with tTAF and PRC1 components, and controls their nucleolar accumulation

What could be the functions of Ctr9t enriched in a distinct compartment of the nucleolus? The characteristic localization of Ctr9t evoked that of tTAF Sa, an essential activator of genes involved in meiotic progression and post-meiotic differentiation of male germ cells [13]. Sa has been reported to be enriched in spermatocyte nucleoli, exhibiting a pattern complementary to Fibrillarin, in addition to its localization to the condensing autosomes [13,39] (Figure 4A). Indeed, Ctr9t and Sa-GFP showed coincident expression in testes and overlapped near perfectly in the spermatocyte nucleolus (Figure 4A). Strikingly, in the absence of *ctr9t*, Sa-GFP failed to maintain the structure in the nucleolus, and abnormally formed smaller speckles scattered in the nucleoplasm (Figure 4B). Sa-GFP signal on autosomes, however, remained unaffected. In addition, Fibrillarin staining confirmed the nucleolus was maintained in *ctr9t* mutants (Figure S4A). The reciprocal analysis revealed that nucleolar localization of Ctr9t remained unchanged in the absence of *sa* (Figure 4C). One of the functions of tTAF Sa is to recruit the subunits forming Polycomb Repression Complex 1 (PRC1) including Polycomb (Pc) to the nucleolus [13]. Consistent with the previous study, a fraction of Pc-GFP was enriched in and around the spermatocyte nucleolus, where we found it partially co-localized with Ctr9t (Figure S4B). In *ctr9t* mutants, Pc-GFP signal in the nucleolus marked with Fibrillarin was dramatically reduced (Figure 4D). Taken together, these results indicate the *ctr9t*-dependent nucleolar enrichment of tTAF Sa and PRC1 component Pc, raising a possibility of functional interaction between Ctr9t, Sa, and Pc in the spermatocyte nucleolus.

**Figure 4.**
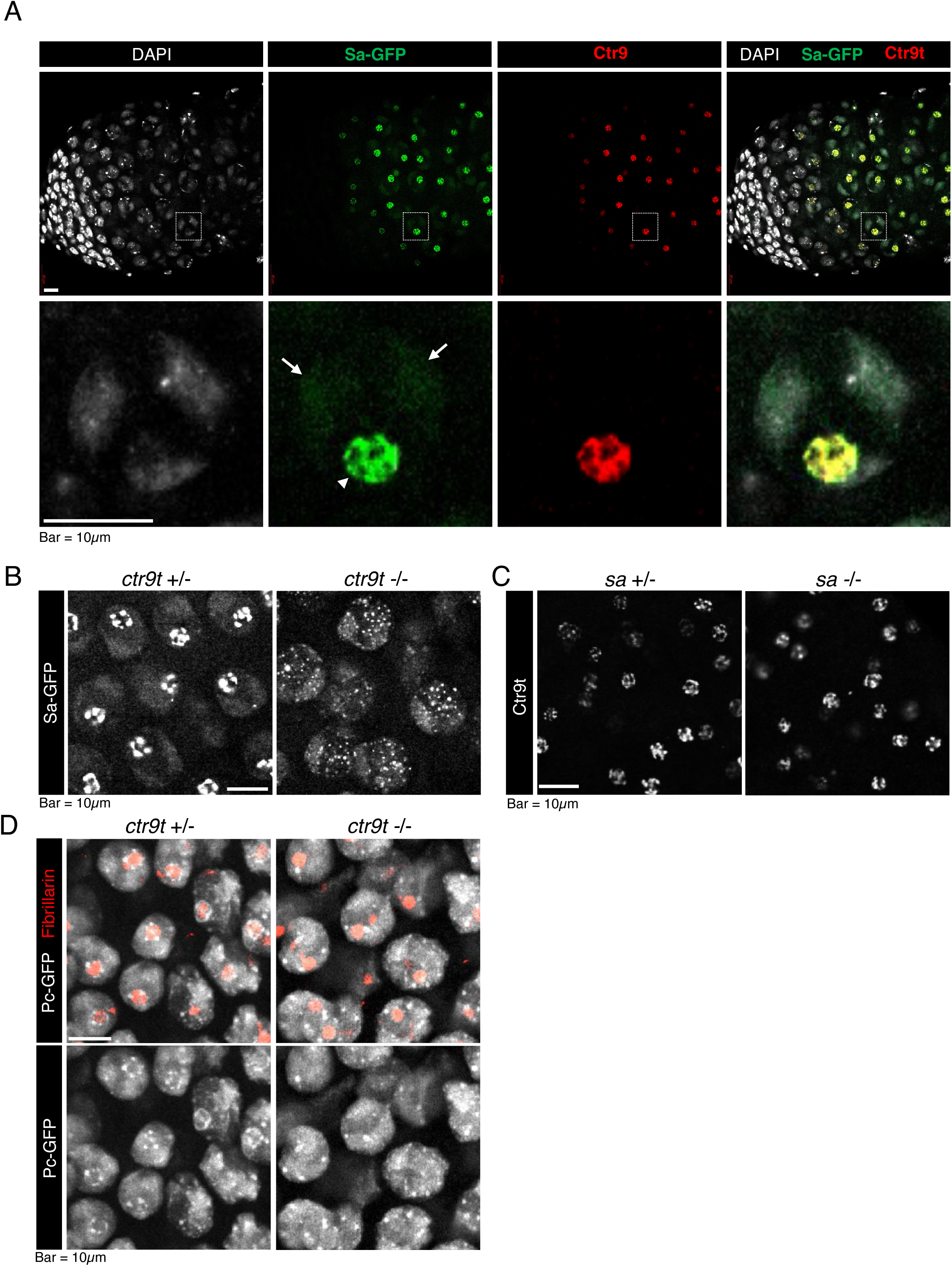
Co-localization and dependency between Ctr9t and tTAF Sa. (A) Testes expressing Sa-GFP (green) immunostained for Ctr9t (red). Magnified images of a spermatocyte nucleus (square box) is shown in the bottom panels. In addition to the signals on autosome regions (arrow), Sa-GFP shows a strong signal enrichment in a nucleolar compartment where Ctr9t predominantly accumulates (arrowhead). (B) Effect of *ctr9t* mutation on the localization of Sa-GFP in spermatocyte nuclei. (C) Effect of *sa* mutation on Ctr9t nucleolar localization. (D) Effect of *ctr9t* mutation on the localization of Pc-GFP in spermacyte nuclei.

### Ctr9t regulates gene expression from sex chromosomes

Germ cells lacking *sa* do not undergo meiosis in testes [11], while those lacking *ctr9t* show clear defects in post-meiotic spermatid differentiation (Figure 2), suggesting the absolute necessity of tTAF Sa, but not of Ctr9t, for meiosis progression. Hence, although Ctr9t is important for maintaining Sa in the nucleolus (Figure 4), it is unlikely that Ctr9t is associated with overall activities of Sa. In order to investigate whether and how Ctr9t acts as a regulator of gene expression, we performed whole testicular transcriptome sequencing, and compared the steady state transcript levels between the control (*KO/CyO* heterozygous and *y w*), *ctr9t* mutants (*KO/KO* and *KO/Df*), and the rescue conditions (*KO/KO* + *bam-Gal4*>UASp-*mTurbo-FLAG-ctr9t*) (Figure 5A). Of the differentially expressed 286 genes in the mutants (p<0.01) (Figure 5B), 163 recovered their expression levels in the rescue condition, and thus were further characterized as *ctr9t*-dependent genes (Table S1) (Figure 5C). Among those, genes regulated by tTAF Sa, such as *mst87F*, *dj*, and *fzo* [39], were not included, and their transcript levels did not change dramatically in *ctr9t* mutants (Figure S5A). Remarkably, of 163 genes, downregulated 91 members contained *kl-2*, *kl-3*, *kl-5*, and *ORY* (*ks-1*) on Y chromosome, all of which are male fertility genes crucial for spermatid differentiation (Figure 5C S5B) [40,41]. The concomitant downregulation of above 4 genes is intriguing given that Y chromosome has only 14 confirmed protein-coding genes (23 including predicted ones among 113 genes totally annotated) [42,43]. Our data also highlighted that X chromosome-located genes occupy more than half of the upregulated members (44 out of 72), and moreover, none of X chromosome genes was included in the downregulated list (Figure 5CD). A more global profiling of all genes expressed in testes (those given transcript fold changes [FC] in differential expression analysis) suggests that the derepression caused by *ctr9t* loss is chromosome-wide (Figure 5EF S5C). For 33 genes expressed from Y chromosome, *ctr9t* loss resulted in various effects on their expression levels (Figure 5E). Nevertheless, we found 15 out of 33 genes showing log2FC>0 located in a specific region (Y:3,258,406-3,365,197) on Y chromosome, implying the effect of *ctr9t* loss depends on the chromosomal topology (Figure 5G S5D). Two more fertility genes on Y chromosome, *WDY (kl-1)* and *CCY (ks-2)* [40,41,44], showed -0.75 and -0.56 of log2FC, and thus were grouped by their *ctr9t*-dependency with *kl-2*, *kl-3*, *kl-5*, and *ORY* (Figure 5G S5D). In contrast, other genes (*FDY*, *Ppr-Y*, and *Pp1-Y2*) dispensable for male fertility did not show *ctr9t*-dependency (log2FC>0). We further confirmed the results of deep-sequencing analysis by RT-qPCR-based measurement (Figure 5H). Altogether, these results suggest that Ctr9t has a role as a regulator of gene expression from sex chromosomes (Figure 6).

**Figure 5.**
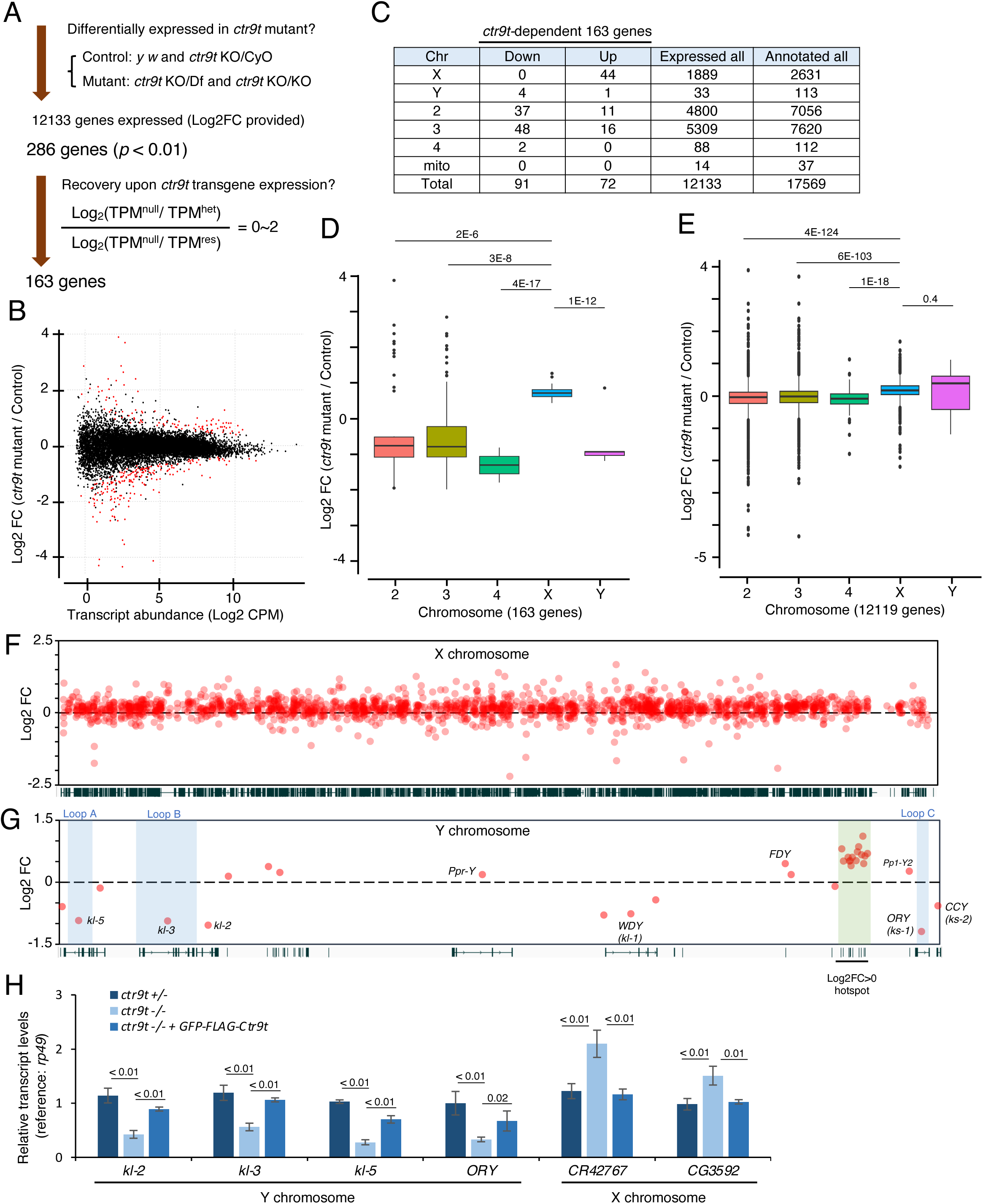
Effect of *ctr9t* mutation on transcript abundance in testes. (A) Analysis scheme of transcriptome sequencing data. Control, mutant, and rescue conditions contain 4 data (2 from *y w* and 2 from *ctr9t^KO/CyO^*), 4 data (2 from *ctr9t^KO/KO^* and 2 from *ctr9t^KO/Df^*), and 2 data from *ctr9t^KO/KO^* + *bam-Gal4*>UASp-*mTurbo-FLAG-ctr9t*, respectively. (B) Volcano plot shows the result of differential expression analysis using EdgeR. Red dot; 286 genes (exact p < 0.01). (C) Table for 163 genes differentially expressed in *ctr9t* mutants and showing expression recovery in rescue condition. Genes were grouped by their locating chromosome and transcript changing patterns (upregulated or downregulated in mutants). (D) Box plot for fold changes of transcript levels with p values (two-tailed unpaired t-test). *ctr9t*-dependent 163 genes are grouped by their locating chromosome. (E) A similar box plot covering all genes extracted from differential expression analysis (n=12119). (F,G) Chromosome-wide landscape of *ctr9t* mutation effect on transcript levels. (H) RT-qPCR measurement of transcript levels for selected genes on sex chromosomes. 4 biological replicates. p values; two-tailed unpaired t-test.

**Figure 6.**
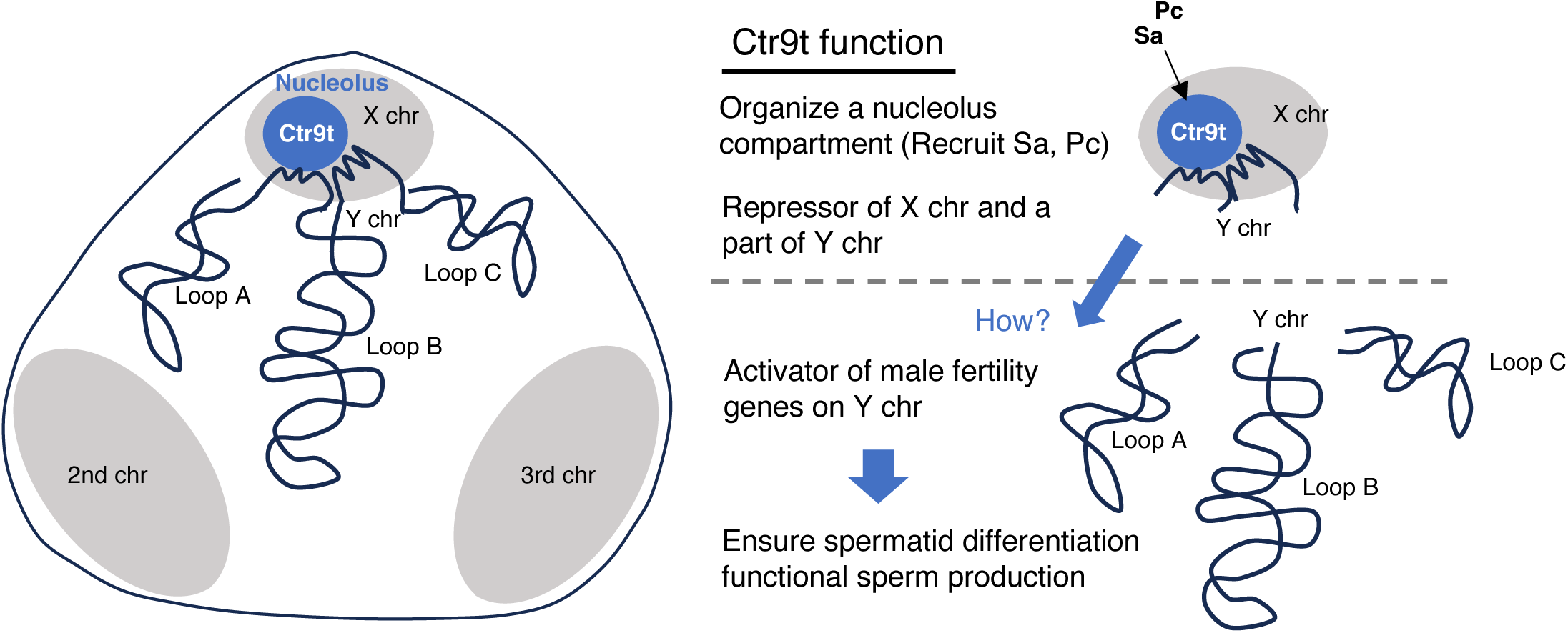
A model for the role of Ctr9t in spermatogenesis. Left; a schematic drawing of *D. melanogaster* spermatocyte nucleus. Condensed X chromosome is universally near the nucleolus. Y loops are formed in the nucleoplasm and thus distant from the nucleolus. Right; a model summarizing the functions of Ctr9t. Ctr9t is exclusively enriched in the spermatocyte nucleolus, required for recruiting tTAF Sa and PRC1 component Pc. Interacting with these and potentially other factors including testis-specific paralogs of Paf1C, Ctr9t regulates gene expression from sex chromosomes. Ctr9t has an important role to activate the set of male fertility genes on Y chromosome, thereby ensuring proper spermatid differentiation and normal male fertility.

## Discussion

Tissue-specific gene regulatory mechanisms can be established by acquiring functionally differentiated paralogs of general transcriptional regulators. This study extended the notion by identifying and functionally characterizing Ctr9t, the testis-specific paralog of Ctr9. Proximity proteome profiling suggests that Ctr9t can act by forming a complex with other paralogs of individual subunits in Paf1C (Figure 2). Nonetheless, further genetic and functional dissection of Paf1t, Cdc73t, Leo1t, and Rtf1t, are necessary to address the possibility. How have Paf1C paralogs been established during evolution? A molecular evolutionary study on tTAFs has shown that individual members arose by independent gene duplication events and evolved rapidly to acquire testis-specific activities through reduced purifying selection, pervasive positive selection, and coevolution [16]. The study suggested that the scenario can be applied to other genes displaying similar patterns of evolution in *Drosophila* lineages and being characterized by testis-enriched expression. Based on the phylogenetic data (Figure 1), we assume that duplication events occurred early in the ancestors of modern *Drosophila* species have contributed to the acquisition of testis-specific paralogs for Paf1C subunits.

Ctr9t is predominantly enriched in a compartment of spermatocyte nucleolus (Figure 3). There, Ctr9t co-localizes near perfectly with a tTAF member, Sa. In addition, nucleolus Sa localization requires *ctr9t* function (Figure 4). Considering these, we assumed that Ctr9t is involved in tTAF-mediated events. However, unlike *sa* mutant germ cells that are unable to undergo meiosis, germ cells lacking *ctr9t* can complete meiosis and progress to the spermatid stages (Figure 2). Moreover, expression of known tTAF targets was not dramatically affected in *ctr9t* mutants (Figure S5). Hence, it would be plausible to conclude that Ctr9t is not associated with overall tTAF activities. In agreement with this, the population of Sa localized in the autosome territories, which is capable of directly controlling tTAF targets, remained unaffected in *ctr9t* mutants [13,39] (Figure 4). What, then, could be the function(s) of Sa in the nucleolus? Perturbation of Sa enrichment in the nucleolus has been reported also in the mutant testes lacking THOC5, a component of THO complex mediating mRNA export [45]. Similar to *sa* mutant, *THOC5* mutant testes show meiotic arrest phenotype. These pieces of evidence highlight the ability of *ctr9t* mutant germ cells to enter post-meiotic stages. Given the fact that nucleolus Sa disruption can be uncoupled from meiotic arrest, potential function(s) of nucleolar Sa may not be necessary for meiosis but rather important for post-meiotic differentiation.

Male fertility genes on Y chromosome, including *kl-2*, *kl-3*, *kl-5*, and *ORY (ks-1)*, depends on Ctr9t function for their expression (Figure 5). We assume this activator role of Ctr9t is relevant for its importance in functional sperm production (Figure 2). However, all of these genes are characteristic by containing gigantic introns, and, accordingly, *kl-3*, *kl-5*, and *ORY* are known to form highly structured co-transcriptional architecture called Y loops in the spermatocyte nuclei [46–50]. Thus, it remains unclear how Ctr9t, which is predominantly enriched in the nucleolus, can activate male fertility genes forming Y loops and thus being distant from the nucleolus (Figure 6). Following the model originally proposed for tTAF Sa [13], sequestering Pc in the nucleolus (Figure 4) may reduce the access of Pc to Y-loops to prevent the silencing of fertility genes. From an evolutionary viewpoint, the set of male fertility genes are maintained by autosome but not by Y chromosome in *D. pseudoobscura,* due to the Y-to-dot translocation event occurred 10-20 million years ago [51,52]. In this regard, the absence of paralogs for *ctr9*, *paf1*, and *leo1* in *D. pseudoobscura* and closely related *D. persimilis* (Figure 1) might imply the particular importance of paralogs for expressing fertility genes from Y loops. Besides the effect on Y chromosome, the mechanism by which Ctr9t causes global repression of X chromosome genes remains elusive. Nevertheless, condensed X chromosome in primary spermatocytes is universally close to the nucleolus, with the located genes being silenced in a chromosome-wide manner by unknown mechanisms [53–55]. In addition, nucleolus is known to contribute to mammalian X chromosome inactivation occuring in female cells [56]. It is therefore possible that Ctr9t directly controls genes on X chromosome. In this sense, the mode of action of Ctr9t could be dissimilar to that of Ctr9, given Paf1C generally acts as transcription activator [22]. Further study on Ctr9t and potentially functional interactors including tTAF Sa and Polycomb (Pc) could provide the mechanistic framework for understanding how X chromosome-wide inactivation occurs in spermatocytes.

Besides serving as platform for ribosome assembly, nucleolus acts as a central hub in mediating multiple events including DNA damage repair, stress response, apoptosis, and chromatin regulation [38,57,58]. Recent studies highlight the interaction of nucleolus with repressive chromatin domains [59,60]. In *Drosophila* spermatocytes, proteins forming Polycomb repressive complex 1 (PRC1) including Pc can have a distinct role in the nucleolus, not being sequestered in an inactive state [13,39,61]. In line with this, the nucleolar region contains elongating RNA polymerase II and TAF1 [15,61]. Ctr9t-dependent enrichment of Pc in nucleolar region implies its involvement in gene repression by Ctr9t (Figure 4). On the other hand, acetylated histones and their testis-specific reader proteins are also enriched in the nucleolus and partially co-localizes with Sa and Pc proteins [62–65]. Hence, spermatocyte nucleolus may exert differential effects on gene expression.

Cytoplasmic dispersion of ectopically expressed Ctr9t in spermatogonia and early spermatocytes (Figure S3) raises a question of what drives the enrichment in the nucleolus. General Ctr9 can be present in the cytoplasm when not forming Paf1C [66]. Perhaps, the interaction between paralogs of Paf1C subunits enables the nucleolar localization. To support this, tPlus3b, one of paralogs of Rtf1 comprising Paf1C, activating *kl-3* and *kl-5* genes similar to Ctr9t, can be found in the nucleolus [33]. Together with previous studies, our work proposes a model whereby different testis-specific gene regulators functionally interact in the nucleolus. Links between paralogs would be of importance in this context. Further studies on *Drosophila* spermatogenesis will reveal otherwise unanticipated functions of the nucleolus in gene regulation.

## Materials and methods

### Fly stocks and cultures

Fly stocks are reared at 25°C on a molasses/yeast medium [5% (w/v) dry yeast, 5% (w/v) corn flower, 2% (w/v) rice bran, 10% (w/v) glucose, 0.7% (w/v) agar, 0.2% (v/v) propionic acid, and 0.05% (v/v) p-hydroxy butyl benzoic acid]. The following stocks were used: *yellow white* (*y w)*, *nos-phiC31*; P{CaryP}attP2 (BL#25710), Df(2R)Exel7177/CyO (BL#7906, deficiency for *ctr9t*), *bam-GAL4/TM6*, *His2Av-mRFP* (BL#23650), *bam^Δ86^*/TM3 (BL#5427), *Sa-GFP* [13], *sa^2^*/TM6B [11], *Pc-GFP* (BL#9593).

### Plasmid construction, fly transformation, mutagenesis

All the primers used for plasmid construction are listed in Table S2. For the establishment of *CG9899/ctr9t^KO^* by CRISPR-Cas9, the left and right homology arms were amplified by pfusion PCR (NEB) using *y w* genomic DNA as template and primer pairs Ti897/898 and Ti899/900. A fragment containing *3xP3-dsRed* marker was amplified using primer pairs Ti673/Mt024 (template: pBlue-cyp40KOarm [67]). These three fragments were cloned into EcoRI-digested pBluescript-II vector using In-Fusion HD cloning kit (Takara). gRNA plasmids were constructed by cloning of annealed oligo DNA (Ti902/903 and Ti904/905) into BbsI-digested pDCC6 vector [35]. The homology arm plasmid and gRNA plasmids were co-injected to *y w* embryos. For UASp-*GFP-3FLAG-ctr9t*, *GFP-3FLAG* and *ctr9t* fragments were amplified by PCR using Ti042/172 (template: pUASp-*GFP-3FLAG-cyp40* [67]) and Ti912/913 (template: testis cDNA), respectively, and cloned into XbaI site of pUASp-K10-attB vector by in-Fusion reaction. For UASp-*mTurbo-FLAG-ctr9t*, *mTurbo* and *FLAG-ctr9t* fragments were amplified by PCR using Ti833/834 (template: UASp-*mTurbo-FLAG-cyp40*, [4]) and Ti835/913 (template: UASp-*GFP-3FLAG-ctr9t*), respectively, and cloned into XbaI site of pUASp-K10-attB vector. The constructs were integrated into *attP2* site.

### Homolog search and phylogenetic analysis

Homologs of individual subunits forming Paf1C were searched by Protein BLAST (BLASTp) using the NCBI platform (https://blast.ncbi.nlm.nih.gov/Blast.cgi). Multiple sequence alignment was performed with full-length polypeptide sequences in MAFFT version 7 (https://mafft.cbrc.jp/alignment/server/index.html) [68]. Phylogenetic tree was created by Neighbor-Joining methods for conserved sites with Jones-Taylor-Thornton (JTT) model, and visualized in phylo.io (https://phylo.io/index.html).

### Antibody generation

A DNA fragment for carboxy-terminal region (position: 790-922) of Ctr9t was amplified by PCR using primers Ti951/952 and cloned into pENTR-D-Topo (Thermo Fisher Scientific). The insert was transferred to pDEST15 destination vector by Gateway LR clonase (Thermo Fisher Scientific). GST-tagged Ctr9t^790-922^ expressed in BL21 (DE3) was purified. Injection of purified antigen to rabbits and guinea pigs, and serum preparation were performed by Kiwa Laboratory Animals Co., Ltd. (https://kwl-a.co.jp/).

### Histochemistry and image acquisition

Adult males were dissected in x1 PBS buffer supplemented with 0.04%(w/v) bovine serum albumin (BSA, Wako), and fixed in 5.3%(v/v) paraformaldehyde (Nacalai) in x0.67 PBS buffer for 10min. The samples were then incubated with 1mM 4’,6-Diamidine-20-phenylindole dihydrochloride (DAPI), Phalloidin Rhodamine X Conjugated (Wako, 1:1000) in PBX buffer (x1 PBS containing 0.2%[v/v] Triton X-100). For immunostaining, fixed testes were washed for 30 min with PBX and incubated with PBX containing 2%(w/v) BSA for blocking. The primary antibody incubation was performed overnight at 4°C, and testes were washed with PBX at 25°C for 1h. The secondary antibody incubation was then performed at 25°C for 2h, and testes were washed with PBX at 25°C for 1h. The following antibodies were used in the indicated dilution; anti-Ctr9t antibody (guinea pig, 1:5000), anti-Ctr9t antibody (rabbit, 1:5000), anti-Vasa antibody (guinea pig, 1:5000) [69], anti-Paf1 antibody (rabbit, 1:200) [24], anti-Cdc73 antibody (rabbit, 1:200) [24], anti-Fibrillarin antibody (Abcam #ab5821, rabbit, 1:500), anti-H3K4me3 antibody (Active motif #61979, mouse, 1:100), anti-guinea pig IgG-Alexa Fluor 488 or 555, anti-mouse IgG-Alexa Fluor 488, and anti-rabbit IgG-Alexa Fluor 488 or 555 (Molecular probes, Invitrogen, 1:500). Images were acquired using LSM900 confocal microscope (Zeiss). Images were processed using Zen (Zeiss) and PowerPoint (Microsoft).

### Immunoblotting

Protein samples were denatured by boiling at 95°C for 3min in protein-loading buffer [2% (w/v) SDS, 100 mM DTT, 0.05% (v/v) BPB, and 10% (v/v) glycerol], resolved by SDS-PAGE, and transferred to 0.2-μm polyvinylidene difluoride membrane (Wako) using the semi-dry system (Trans-blot Turbo, Bio-Rad). The membrane was blocked in 4% (w/v) skim milk (Nacalai) in 1x phosphate-buffered saline (PBS) supplemented with 0.1% (v/v) Tween 20 and further incubated with the anti-Ctr9t antibodies (1:1000; guinea pig or rabbit). Coomassie brilliant blue (CBB) staining serves as protein loading control. Images were processed using Fiji.

### Proximity proteome using TurboID

mini(m)Turbo fusion proteins were expressed in germ cells under *bam* promoter activity using Gal4/UASp system. After eclosion, male progenies were reared at 25°C for 3 days in the modified molasses/yeast medium supplemented with 100 μM biotin (Nacalai). Biotinylated proteins were purified from 200 testes as described [4], and resolved by SDS-polyacrylamide gel electrophoresis (SDS-PAGE) in 5 to 20% precast gel (ATTO). Proteins in gel particles were digested with trypsin, and analyzed by liquid chromatography tandem MS (LC_MS/MS) using Q Exactive and UltiMate 3000 Nano LC (Thermo Fisher Scientific) in CoMiT Omics Center (Osaka University, Japan). The mass spectrum data were analyzed by Mascot v2.5.1 (Matrix Science).

### Transcriptome analysis

From 150 to 200 testes of ≤3-day old adults, total RNAs were extracted using TRIzol LS reagent (Thermo Fisher Scientific) following the manufacturer’s protocol and precipitated in the presence of 50% (v/v) isopropanol and 20μg of glycogen (Nacalai) overnight at −20°C. After centrifugation (20,000g, 20 min, 4C), the pellet was rinsed twice with 80% (v/v) ethanol and then resuspended in RNase-free water. After treatment with TURBO DNase (Thermo Fisher Scientific), RNA mixture was added with equal volume of phenol:chloroform:isoamyl alcohol (25:24:1, pH 5.2, Nacalai), vortexed, and centrifuged (20,000g, 3min), and the supernatant was collected. After repeating this step, RNA in the supernatant was precipitated in the presence of 0.3 M NaOAc (pH 5.2) and 75% (v/v) ethanol overnight at −20°C. After centrifugation (20,000g, 20min, 4°C), the pellet was rinsed twice with 80% (v/v) ethanol, resuspended in RNase-free water, and stored at −80°C until shipment. Library construction and sequencing were performed by Rhelixa (Japan). After enriching polyadenylated RNA using NEBNext Poly(A) mRNA Magnetic Isolation Module, libraries were made by using the NEBNext Ultra II Directional RNA Library Prep Kit. Sequencing was performed in NovaSeq 6000 (Illumina). Paired reads (150+150nt) were mapped on *D. melanogaster* genome (Drosophila_melanogaster.BDGP6.32.dna.toplevel.fa, https://ftp.ensembl.org/) by STAR (version 2.7.10b, --outFilterMultimapNmax 100 --outSAMmultNmax 1 -- outMultimapperOrder Random). Conversion to binary alignment map (bam) format and sorting were done using Samtools. Gene exon mappers were counted by featureCounts (version 2.0.1, -M -p -a dmel-all-r6.32.gtf (https://ftp.flybase.net/)). Using counts on exons of genes, Transcripts Per Kilobase Milliion (TPM) was measured. Bedgraph files were generated by bedtools genomecov (-bga -split -scale). Using the scale option, reads per million (RPM) in total exon mappers were obtained. After sorting (sort -k1,1 -k2,2n), bedgraph files between biological replicates were combined by bedtools unionbedg. Mean RPM in counting columns were given by awk command. Final bedgraph was visualized in IGV (https://software.broadinstitute.org/software/igv/). Using the exon read counts on genes provided by featureCounts as input, the differential expression analysis was conducted by edgeR using R version 4.2.3, between control (4 replicates; 2 from *y w* and 2 from *ctr9t*^KO/CyO^) and mutant (4 replicates; 2 from *ctr9t*^KO/Df^ and 2 from *ctr9t*^KO/KO^) conditions. Lowly expressed genes were filtered out with filterByExpr (12149 genes remained, 12133 genes had information of chromosome location). Differentially expressed genes were identified with the threshold of exact *p* < 0.01 (n=286). Of note, only 56 genes were extracted when FDR<0.05 was used as threshold. For the 286 genes, expression recovery in the rescue condition (2 replicates from *ctr9t*^KO/KO^ + *mTurbo-FLAG-ctr9t*) was judged when the values of Log_2_(TPM^mutant^/TPM^control^)/Log_2_(TPM^mutant^/TPM^rescue^) ranges from 0 to 2. To generate Figure 1C, raw data were collected from PRJEB22205. Mapping and counting were done as described above but in a single-end mode. Since featurecounts failed to count mappers on *Leo1/Atu* (probably due to the presence of overlapping non-coding RNA gene), *Leo1/Atu* mappers were separately counted with samtools view.

### Quantitative reverse transcription PCR

RNA was extracted from ∼10 testes of ≤3-day-old flies for each condition using TRIzol LS reagent (Thermo Fisher Scientific) following the manufacturer’s protocol. Using DNase I (NEB)–treated RNA, cDNA was synthesized with 2.5 μM oligo(dT) adaptor using SuperScript III reverse transcriptase (Thermo Fisher Scientific). Quantivative reverse transcription PCR (qPCR) reaction was performed using KAPA SYBR FAST qPCR Master Mix (Roche) and gene-specific primers (Table S2) in QuantStudio 5 Real-Time PCR system (ABI). Relative transcript levels were calcularated by 2^-𝛥𝛥Ct^ method using *rp49* as reference.

### Male fertility test

Single male was mated with six *y w* virgin females for the first 3 days and with another six *y w* virgin females the following 2 days at 25°C. The total number of hatched eggs was counted.

## Supplementary figure legends

**Figure S1.**
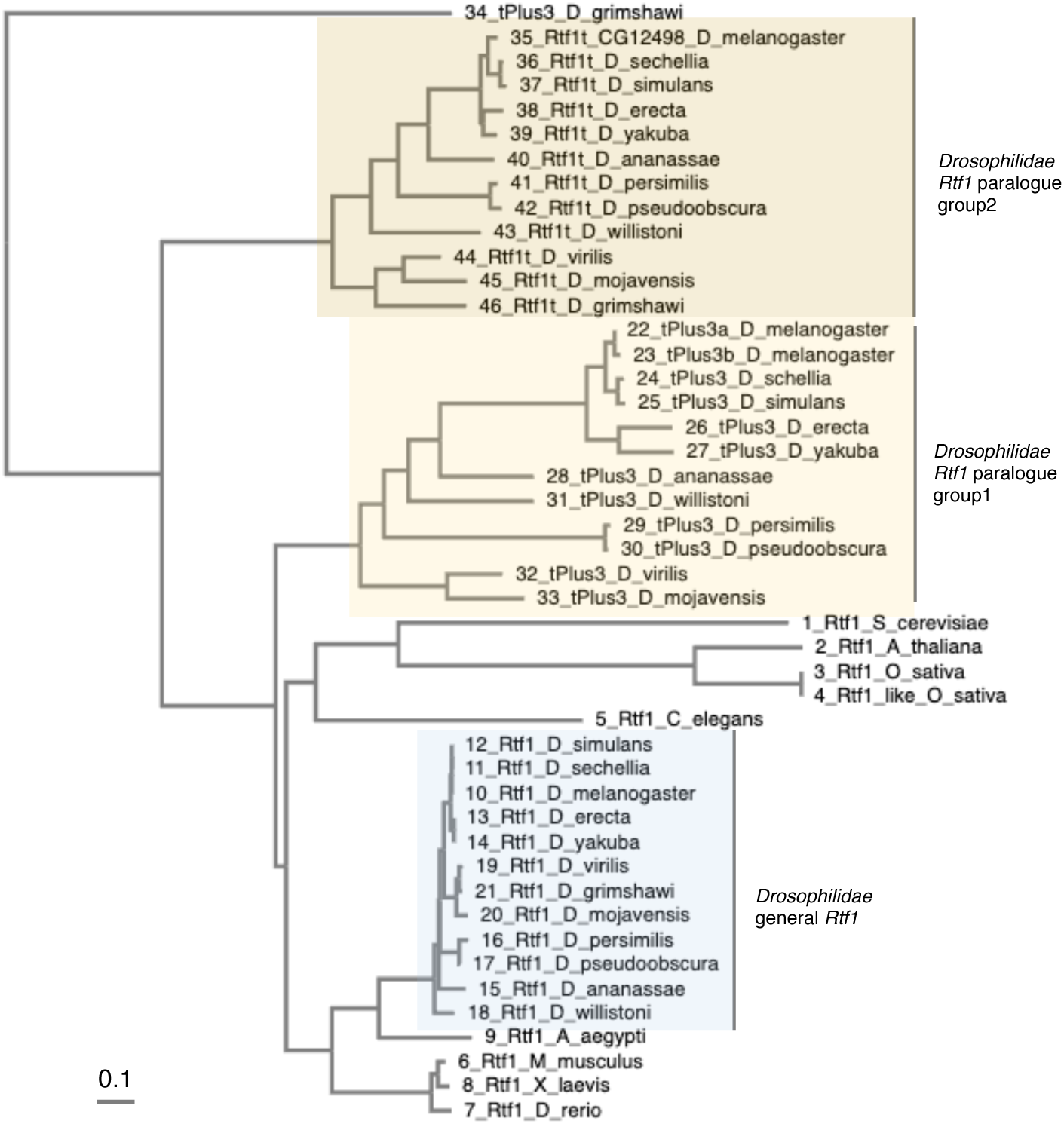
Phylogenetic analysis of Rtf1 homologs. Multiple alignment and phylogenetic analysis were done with Rtf1 protein sequences. Branch lengths measure the expected substitutions per site as indicated in the scale bar.

**Figure S2.**
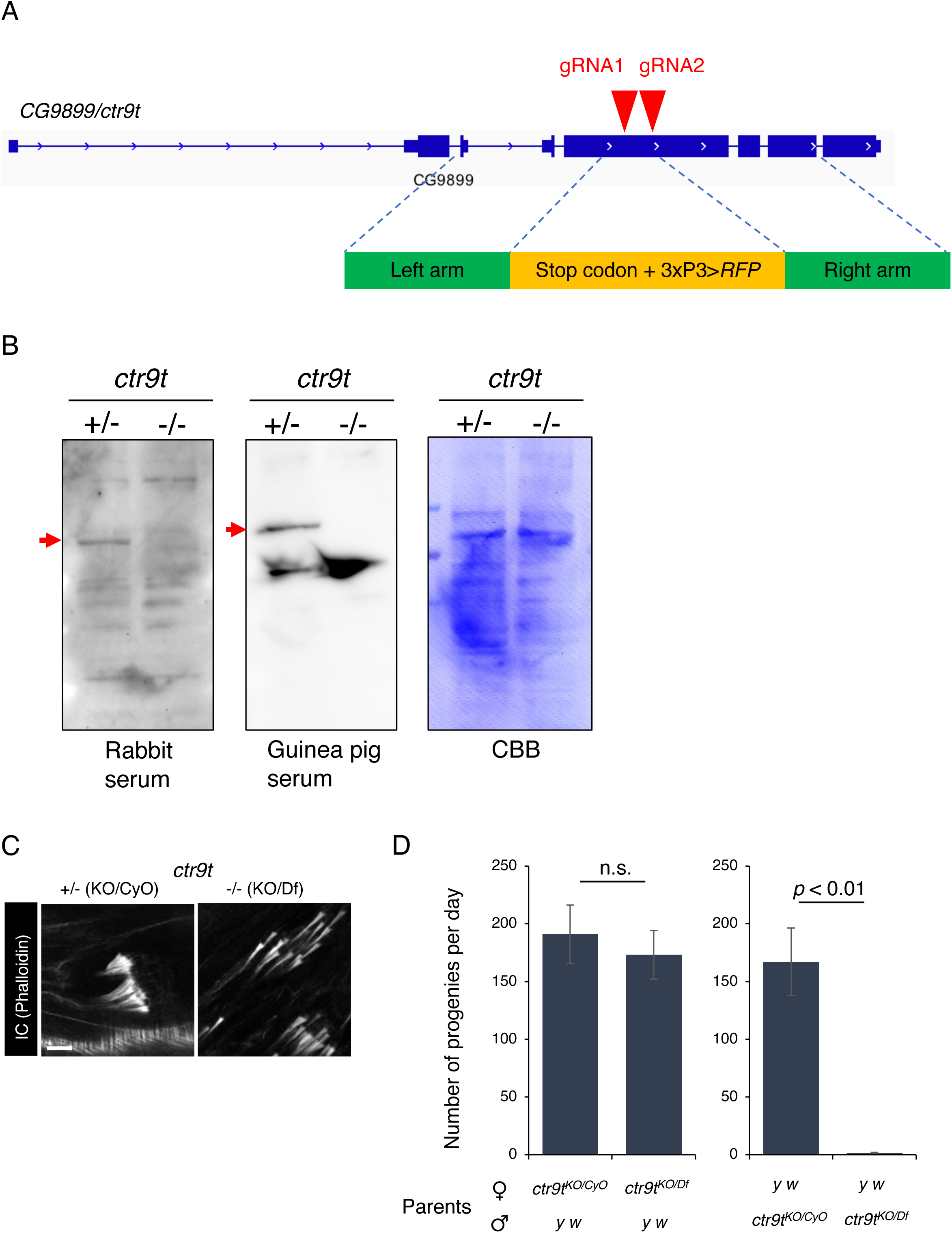
Construction and analysis of *ctr9t* null allele. (A) CRISPR-Cas9-based genome editing for the generation of *ctr9t* null mutant. Guide RNA position and homology arm regions were indicated. (B) Immunoblotting using anti-Ctr9t antibodies raised in rabbit or guinea pig. Lysates from *ctr9t* heterozygous (+/-) and homozygous mutant (-/-) testes were compared. CBB staining serves as protein loading control. (C) IC organization in *ctr9t* transheterozygous mutant spermatids (-/-, KO/Df). Control; sibling heterozygous (+/-, KO/CyO). (D) Female and male fertility of *ctr9t* transheterozygous mutants (-/-, KO/Df) compared to the sibling heterozygous control (+/-, KO/CyO).

**Figure S3.**
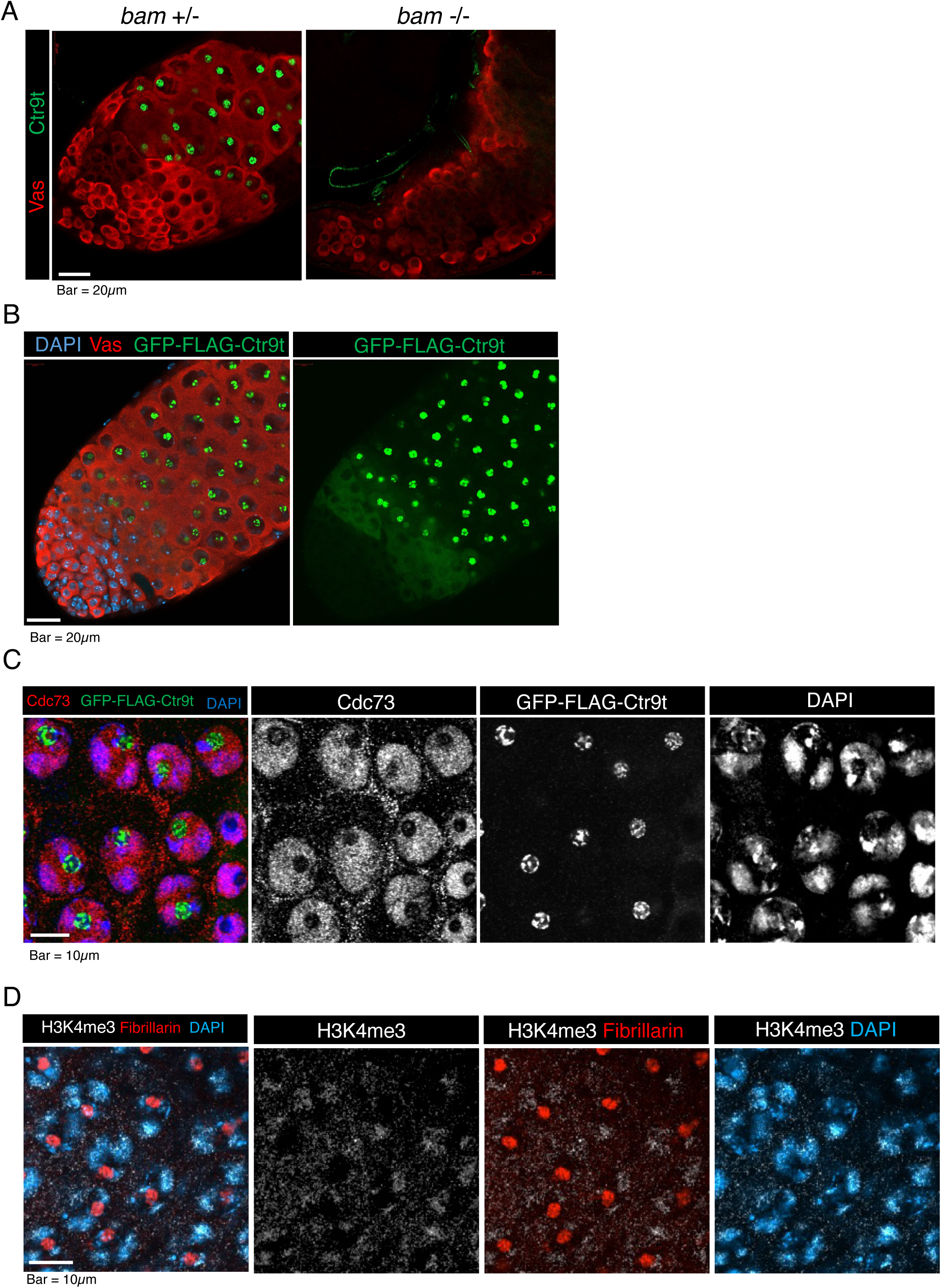
Expression and localization of Ctr9t. (A) Immunostaining of Ctr9t (Green) in the presence or absence of *bam*. +/-; *Δ86/TM3*, -/-; *Δ86/Δ86*. Germ cells (Vas, red). (B) Fluorescence signals from GFP-FLAG-Ctr9t (Green) expressed under the control of *bam* promoter activity. Germ cells (Vas, red) and DNA (DAPI, blue). (C) Immunostaining of general Paf1C subunit, Cdc73, in spermatocytes expressing GFP-FLAG-Ctr9t. (D) Immunostaining of H3K4me3 (white) and Fibrillarin (red) in spermatocytes. DNA (DAPI, blue). H3K4me3 signals overlap with autosomes but under detectable level in the nucleolar region.

**Figure S4.**
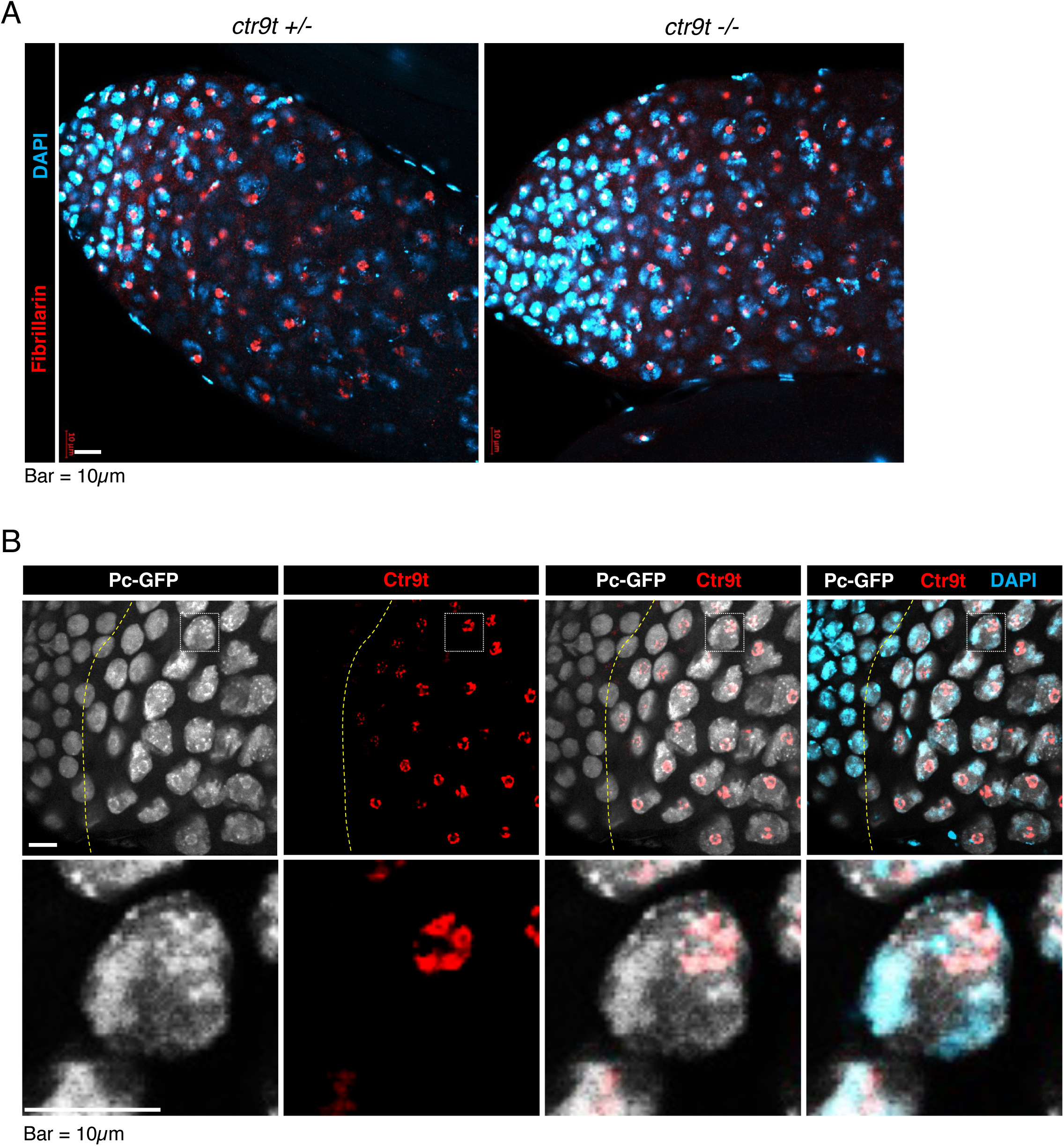
Nucleoli in testes lacking *ctr9t*, Pc-GFP localization in testes. (A) Nucleolus (Fibrillarin, red) and DNA (DAPI, blue) in the apical end of testes of *ctr9t* heterozygous control (+/-, KO/CyO) and homozygous mutant (-/-, KO/KO). (B) Testes expressing Pc-GFP (white) immunostained for Ctr9t (red). DNA (DAPI, blue). Dotted yellow line roughly delineates the boundary between spermatogonia and spermatocytes. Nucleolar enrichment of Pc-GFP coincided with the expression of Ctr9t. Magnified images of a spermatocyte nucleus (square box) was shown in the bottom panels.

**Figure S5.**
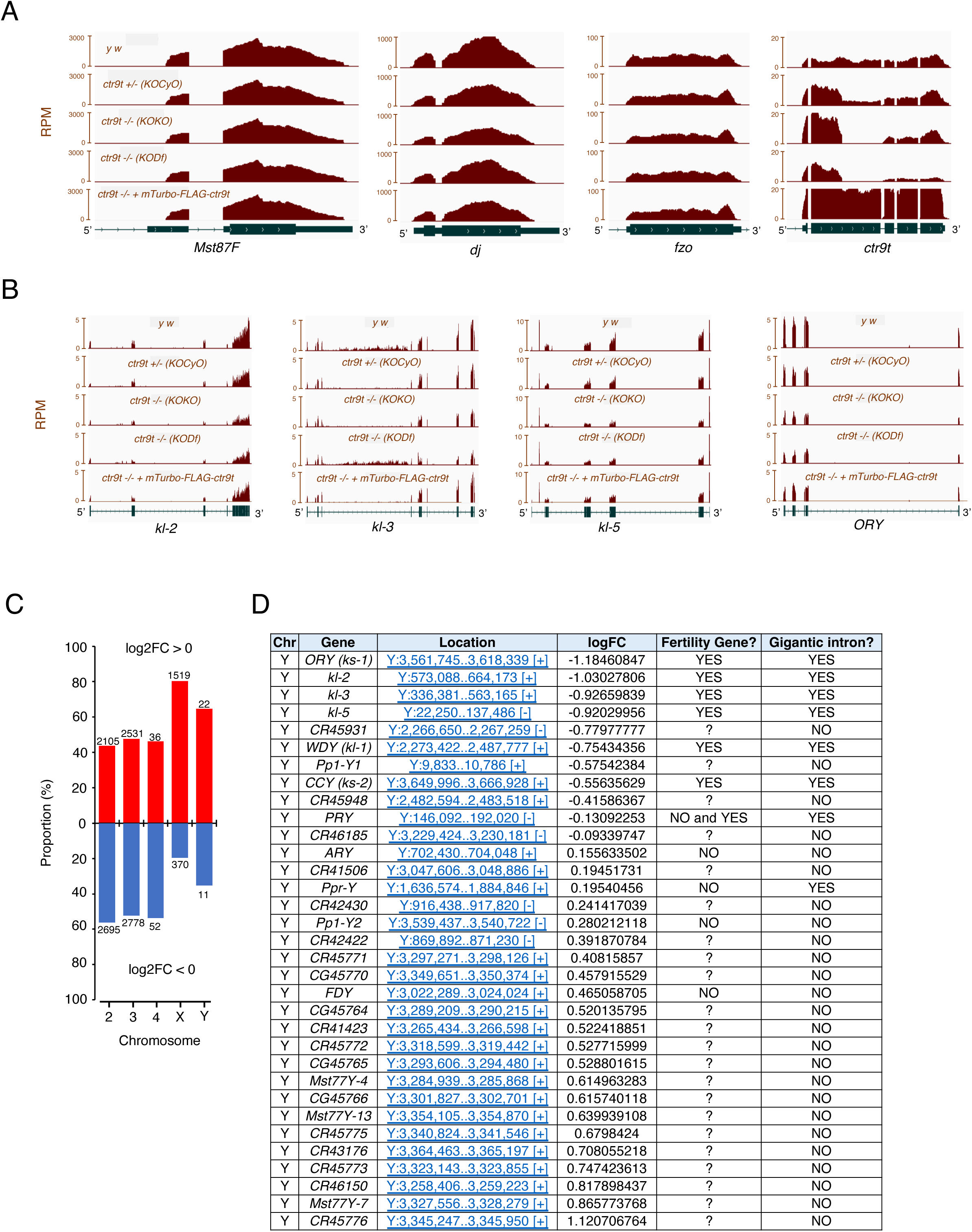
Effect of *ctr9t* loss on testicular transcriptome. (A) Bedgraph shows RPM of known targets of tTAF (*Mst87F, dj, fzo*) in control (*yw* and *ctr9t*^KO^/CyO), mutant (*ctr9t*^KO/KO^ and *ctr9t*^KO^/Df), and rescue conditions (*ctr9t*^KO/KO^ + mTurbo-FLAG-Ctr9t). Bedgraph for *ctr9t* is also provided to confirm the analyzed conditions. (B) Bedgraph shows RPM of Y chromosome-located male fertility genes. (C) Bar graph summarizes the numbers of genes having positive or negative values of transcript log2FC (*ctr9t* mutant compared to control) on each chromosome. (D) List of 33 genes expressed from Y chromosome (those given log2FC values after differential expression analysis).

## Acknowledgement

We thank Dr. Prof. John Lis and Dr. Prof. Margaret Fuller for reagents. We also thank Minori Kawaguchi for her help with preparation of Ctr9t antigens. We thank Chisato Yanagisawa, Momiji Hirayama, and Dr. Rizky Mutiara for their helps with maintenance of fly lines. We are grateful to Bloomington Drosophila Stock Center for providing fly stocks. We also thank all members in our lab for their insightful discussion and suggestions.

## Author contributions

Conceptualization: TI

Methodology: TI and TK

Investigation: TI, JZ, and TK

Supervisions: TI and TK

Writing-original draft: TI

Writing-review and editing: TI and TK

## Data availability

Newly generated transcriptome data are available with BioProject accession ID: PRJNA1094673.

